# Dominant gingers – discovery and inheritance of a new shell polymorphism in the great pond snail *Lymnaea stagnalis*

**DOI:** 10.1101/2023.01.24.525303

**Authors:** Matthijs Ledder, Yumi Nakadera, Alexandra Staikou, Joris M. Koene

**Affiliations:** Ecology and Evolution, A-LIFE, Vrije Universiteit Amsterdam, the Netherlands; Department of Zoology, School of Biology, Aristotle University of Thessaloniki, Greece

**Keywords:** Mendelian inheritance, genetics, evolution, life history, simultaneous hermaphrodites

## Abstract

Color polymorphism is a classic study system for evolutionary genetics. One of the most color-polymorphic animal taxa is mollusks, but the investigation for the genetic basis of color determination is often hindered by their life history and limited availability of genetic resources. Here we report on the discovery of shell color polymorphism in a much-used model species, the great pond snail *Lymnaea stagnalis*. While their shell is usually beige, some individuals from a Greek population show a distinct red shell color, which we nicknamed Ginger. Moreover, we found that the inheritance fits simple, single-locus Mendelian inheritance with dominance of the Ginger allele. We also compared crucial life history traits between Ginger and wild type individuals, and found no differences between morphs. We conclude that the relative simplicity of this polymorphism will provide new opportunities for a deeper understanding of the genetic basis of shell color polymorphism and its evolutionary origin.

## Introduction

Visible polymorphisms are valuable to expand the understanding of underlying genetics as well as evolutionary processes involved. The former is well demonstrated by the seminal work of Mendel (1865), providing the foundation of genetics. The latter can inform us about the maintenance of genetic variation and changes in selective pressure in natural populations, as in the case of industrial melanism in the peppered moth *Biston betularia* (reviewed in Cook 2003; Cook and Saccheri 2013). There are a vast number of studies which demonstrate the significance of color polymorphism and aided us in expanding the understanding of mutation and selection (e.g., Valverde et al. 1995; Quattrocchio et al. 1999; Nadeau et al. 2016; Kellenberger et al. 2019; Feiner et al. 2022).

Besides fundamental insights, the aesthetic appeal of color polymorphism is highly compatible with citizen science, providing outreach opportunities and at the same time advancing the research with massive amounts of data (e.g., Silvertown et al. 2011; Kerstes et al. 2019). For example, the grove snail *Cepaea nemoralis* (and the closely related *C. hortensis*) is a historical model species of shell color polymorphism, given their various ground colors (yellow, pink and brown), banding pattern (up to five bands) as well as band and lip colors. Previous studies demonstrated that these phenotypes are controlled by a series of eight or more genes (e.g., Gonzalez et al. 2019; Kerkvliet et al. 2017). Recently, citizen science provided massive data to confirm that *Cepaea’s* shell morphs are correlated with their habitat, e.g., urban snails tend to be more often yellow than pink, potentially due to thermal selection (Cain and Sheppard 1950, 1954; Kerstes et al. 2019, but see also Silvertown et al. 2011).

The Gastropoda is one of the most color polymorphic animal groups (e.g., Lee et al. 2022; Luttikhuizen and Drent 2008; Liu et al. 2009, reviewed in Williams 2017; Gefaell et al. 2022). However, except for a few cases like *Cepaea*, the genetic basis, evolution and ecological meanings of these colors largely remain to be examined. The major logistical hinderances stem from the life histories of many gastropod species, such as their long, slow life cycle or the presence of a planktonic life stage. Even more crucially, it is challenging to obtain progenies from desired crosses, due to e.g., low mating rate or fecundity. Consequently, their life history limits the genomic insights of color polymorphism in other mollusks than *Cepaea*.

We here report the discovery of a new shell polymorphism in a Greek population of *Lymnaea stagnalis*, potentially a new model to expand the understanding of color polymorphism at the genetic, ecological and evolutionary level. We found that some snails from this field population have a distinctly red shell color (nicknamed Ginger), while the shell color of this species is usually beige (Fig. 1B). To see whether the Ginger snails could provide a new model system for studying shell color polymorphism, we tested (1) if Ginger snails show any difference in life history traits compared to the wild-type snails from the same population and (2) how this shell trait inherits to the next generation.

**Fig. 1.**
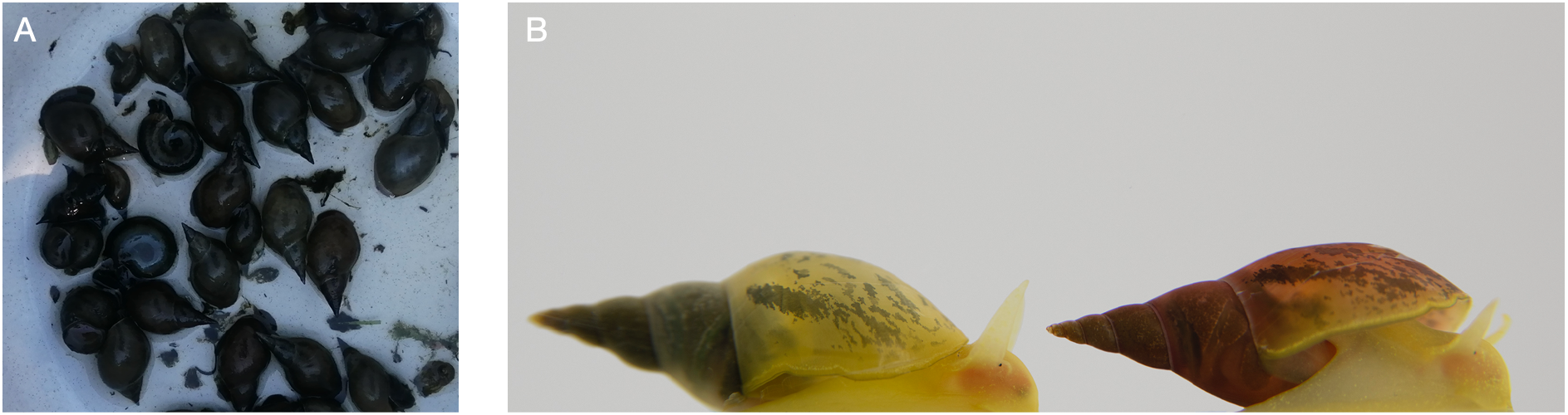
(A) Freshly collected *L. stagnalis* and other freshwater snail species from Kerkini lake, Greece. © Alexandra Staikou. (B) The lab-reared Wild Type (WT) and Ginger (G) snails. © Cathy Levesque.

## Material and Methods

*L. stagnalis* is a hermaphroditic freshwater pulmonate, commonly distributed across the globe (Fodor et al., 2020; Kopp et al. 2012). This species is a simultaneous hermaphrodite, possessing functional organs for both sexes, but when they copulate, one snail acts as sperm donor (male role) inseminating a recipient snail (female role, i.e. unilateral mating); roles can be swapped immediately after this interaction. Moreover, they are capable of self-fertilization (selfing) without showing obvious effects of inbreeding depression (e.g., Coutellec and Lagadic, 2006; Koene et al., 2008; Escobar et al. 2011).

We collected the snails from the shallow waters in Kerkini lake, Central Macedonia, Greece (GPS coordinates 41.24979 23.20994). There was little to no water flow and the water depth was 30 to 50 cm at the site of collection. We collected the snails in July 2021 and transported the snails to Vrije Universiteit Amsterdam for further investigation.

We used 64 age-synchronized snails from the F2 generation of this Greek population (28 Ginger (G), 36 wild type (WT)). In the lab, we kept the snails in a flow-through tank with low-copper water at 20 ± 1 °C. The light:dark cycle was 12:12h. To keep track of mating history of each snail, we raised these snails as virgins. That is, well before their maturation (Shell length ca. 1.5 cm), we placed and raised each snail in an individual container until and beyond their maturation. These containers were perforated (ca. 460 ml) and placed in a large flow-through tank, providing the snails with clean aerated water while they could not have physical contacts with others. We replaced the containers every two weeks, fed the snails with broad leaf lettuce ad libitum, and occasionally provided fish food (TetraPhyll®) as supplemental food.

### Life history comparison

We compared their onset of egg laying, body size, fecundity and mating behavior between G and WT. We started the monitoring when the snails were ca. two-month-old. To record their onset of egg laying, we checked for the presence of eggs in containers every week for 10 weeks. We define the onset of egg laying as the week when an individual produced their first eggs via self-fertilization. To compare the body size between morphs, we measured their shell length as the proxy of body size, using a Vernier caliper (DialMax, 0.1 mm) at Week 7 from the start of our monitoring. To examine their fecundity, at week 9, we provided the snails each with a clean container to enhance egg laying (Ter Maat et al.,1983). After two days, we collected all the egg masses laid to scan them using a flatbed scanner (Canon LiDE 220, van Iersel et al., 2014). Using these images, we counted the number of eggs using ImageJ (ver1.53t, Schneider et al., 2012).

To compare mating behavior of G and WT from the Greek population, we let these snails mate with the snails from our standard laboratory strain maintained at Vrije Universiteit Amsterdam, which never showed the Ginger phenotype since the start of the breeding facility in the 1960s. We selected 12 G and 12 WT for mating observations. First, we isolated 24 lab strain snails for two days, to standardize their mating history. In Week 11, we observed their matings, following our standard protocol (e.g., Nakadera et al. 2015). In short, we placed one Greek and one lab snail into a container, and every 15 min recorded for each pair the following behavioral categories: no contact, shell mounting, probing with the preputium and inseminating (e.g., Koene, 2010). Based on this record, we determined mating rate (the number of mated individuals) as well as the duration of a) mating latency (how long they took to start courtship, i.e. shell mounting), b) courtship (the duration between shell mounting and probing) and c) insemination.

### Inheritance of shell color

To reveal the mode of inheritance, we carried out two experiments. First, we counted the number of G or WT progenies from self-fertilized egg masses. Second, we crossed the Greek snails with the snails from our standard lab strain to measure the change of phenotypic ratios in progeny.

Selfing allowed us to see the mode of inheritance in this shell color polymorphism, without having to take the genotype of the mating partners into account. Note that selfing does not produce genetic clones, but the offspring are still produced via the fusion of a sperm and egg of the same individual within its reproductive tract. To obtain selfed progenies from G and WT, we reared them in isolation (i.e., they were virgin) and collected one egg mass per capita (N: G = 15 out of 21, WT = 21 out of 36). Then, we incubated these egg masses in small plastic vials (ca. 10 ml) with water. After hatching, we kept rearing them in non-perforated containers placed in a flow-through tank, until we were able to see the shell color of all hatchlings clearly. Then, we counted the number of offspring with G or WT phenotypes for each parent.

Letting the Greek snails cross-fertilize with the lab snails allowed us to verify the mode of inheritance predicted from the previous selfing test, since we knew that the lab snails did not have a Ginger allele. To obtain outcrossed progenies, we let the Greek snails mate with the lab snails (see above). After observing the initial matings, we kept these pairs together for three days during which they could mate several times with the same partner. Next, we removed the lab snails and provided new containers. After five days, we collected the outcrossed egg masses from the Greek snails. We reared these outcrossed eggs and hatchlings using the same method as explained above, until their shell color was clearly visible. Lastly, we counted the number of offspring with G or WT phenotypes for each parent.

### Statistics

We conducted all the analyses in R (ver. 4.2.1, R Core Team, 2022). To compare the life history traits between morphs, we mostly used *t*-tests, except mating rate (Chi-square test) and mating behavior data (Wilcoxon tests). To examine if the phenotypic ratio of offspring from heterozygous G mothers deviated from the expectation based on Mendelian single-locus inheritance, we used one sample *t*-tests with the expected mean of 0.75 or 0.5.

## Results

### General observation of Ginger snails

The shell of *L. stagnalis* usually is beige to slightly yellow, but the Ginger snails showed a distinct pink to red color on their shells (Fig. 1B). Both types of shells are relatively transparent, so that we can partly see coloration and patterns of soft body. In the field, their shells are typically covered with algae and fungus and their soft body is much darker than the lab reared snails (Fig. 1A), likely due to their food and higher UV exposure (Ahlgren et al. 2013). Thus, the shell color differences are almost unnoticeable in field collected snails. In fact, we only noticed the polymorphism in the F1 generation reared in the lab. In retrospect, we also observed that the color is present in shell specimens after the snail died. We did not detect any snails with intermediate shell colors. Lastly, although we have tried using several optic methods during their development, these shell colors become clearly distinguishable only two months after hatching (Shell length: ca. 1 cm).

### Life history comparison

We did not detect any difference in life history traits between G and WT of the Greek population (Fig. S1). There was no difference in the onset of egg laying (*t*_47.78_ = −0.16, *P* = 0.869), body size (*t*_52.16_ = 0.60, *P* = 0.548), fecundity (*t*_34.03_ = −0.58, *P* = 0.562) between G and WT. Regarding mating behavior, when the Greek snails mated with the lab snails, the mating rates were not different between morphs (χ^2^1 = 2.27, *P* = 0.132). When they mate, all Greek snails acted as female first. In mating pairs, we did not see any difference in mating behavior between morphs (mating latency: *W* = 38.5, *P* = 0.673, courtship: *W* = 41, *P* = 0.827, insemination: *W* = 48, *P* = 0.745, Fig. S1).

### Inheritance of shell color

When we monitored the phenotype of self-fertilized offspring, we did not find any G offspring produced by WT mothers, suggesting that the G allele is dominant and the WT are recessive homozygous for this locus (Fig. 2). Also, four out of 15 G mothers produced only G offspring, suggesting their homozygosity (Fig. 2). The rest of G mothers produced a few WT offspring, and the ratio of G produced did not significantly deviate from 0.75, as predicted in the model of single-locus Mendelian inheritance (Mean = 0.716, 95% CI: 0.652 - 0.781, *t*_10_ = - 1.17, *P* = 0.271).

**Fig. 2.**
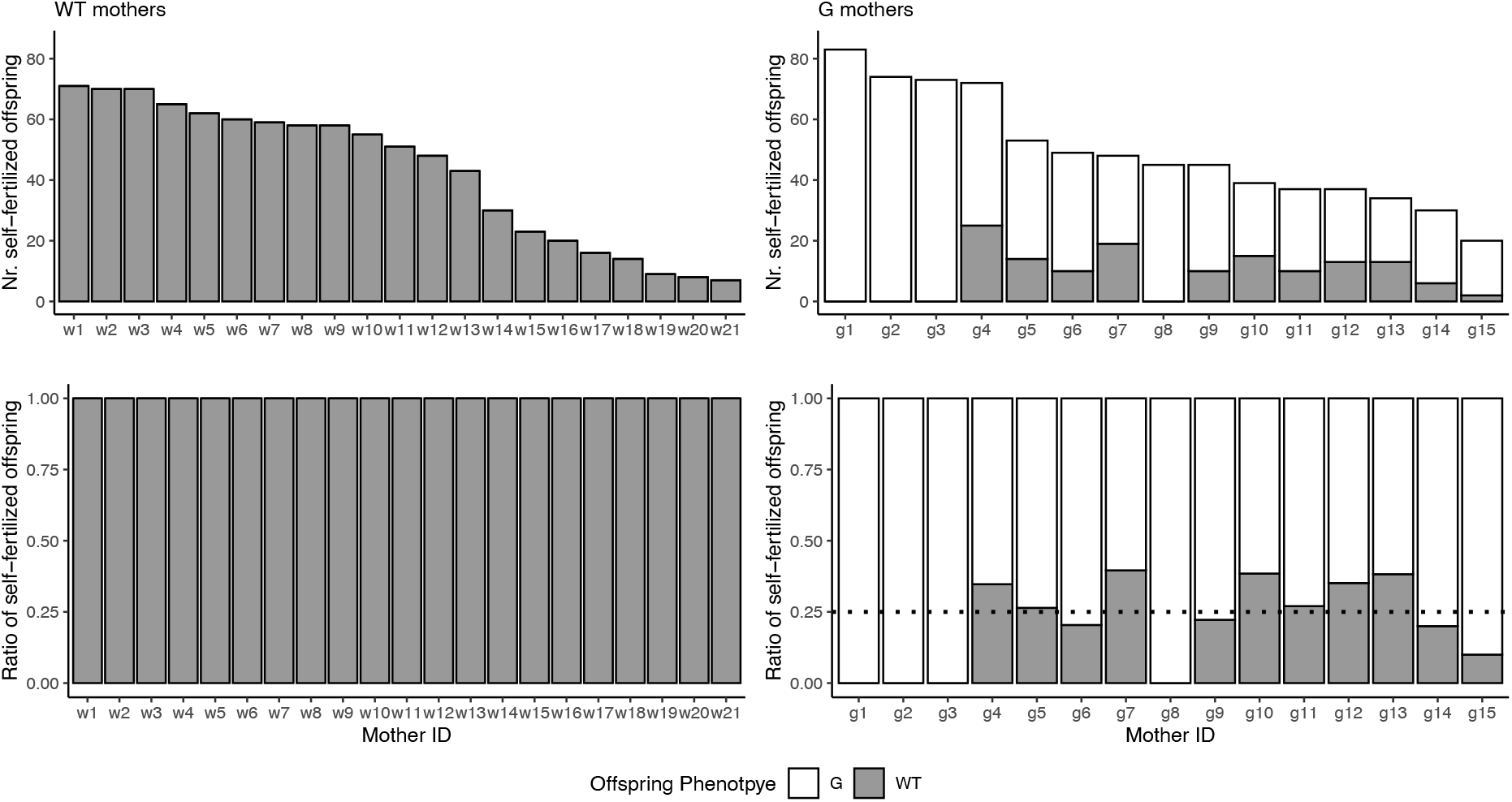
The phenotype of selfed offspring from WT and G mothers. The top panels show the number of offspring, and the bottom ones show the ratios. The individuals are ordered by the total number of offspring we counted (i.e. the used mother IDs are a result of this and are non-informative). The dotted line in the right bottom panel indicates the expected ratio of segregation for single-locus Mendelian inheritance from heterozygous mothers.

After crossing with the lab strain snails, WT mothers from the Greek population again did not produce any G offspring (Fig. 3). For G mothers, we saw that two individuals produced only G offspring, suggesting that they were homozygous, producing heterozygous G offspring (Fig. 3). The rest of G mothers produced both phenotypes, and the ratio of G offspring significantly deviated from 0.75 (Mean = 0.630, 95% CI: 0.537 - 0.724, *t*_9_ = −2.88, *P* = 0.018), indicating that the segregation rate altered after mating with the lab snails. However, the segregation ratio was also significantly deviated from 0.5 as expected in Mendelian inheritance model (*t*_9_ = 3.14, *P* = 0.012). Moreover, the dataset of outcrossed offspring included six G mothers for which we also measured the ratio of selfed offspring (Fig. S2). This confirms that two homozygous G mothers produces only G offspring, both through self-fertilization and outcrossing. Two heterozygous G mothers increased the ratio of WT offspring after mating, while two other heterozygous G mothers did not show such a pattern.

**Fig. 3.**
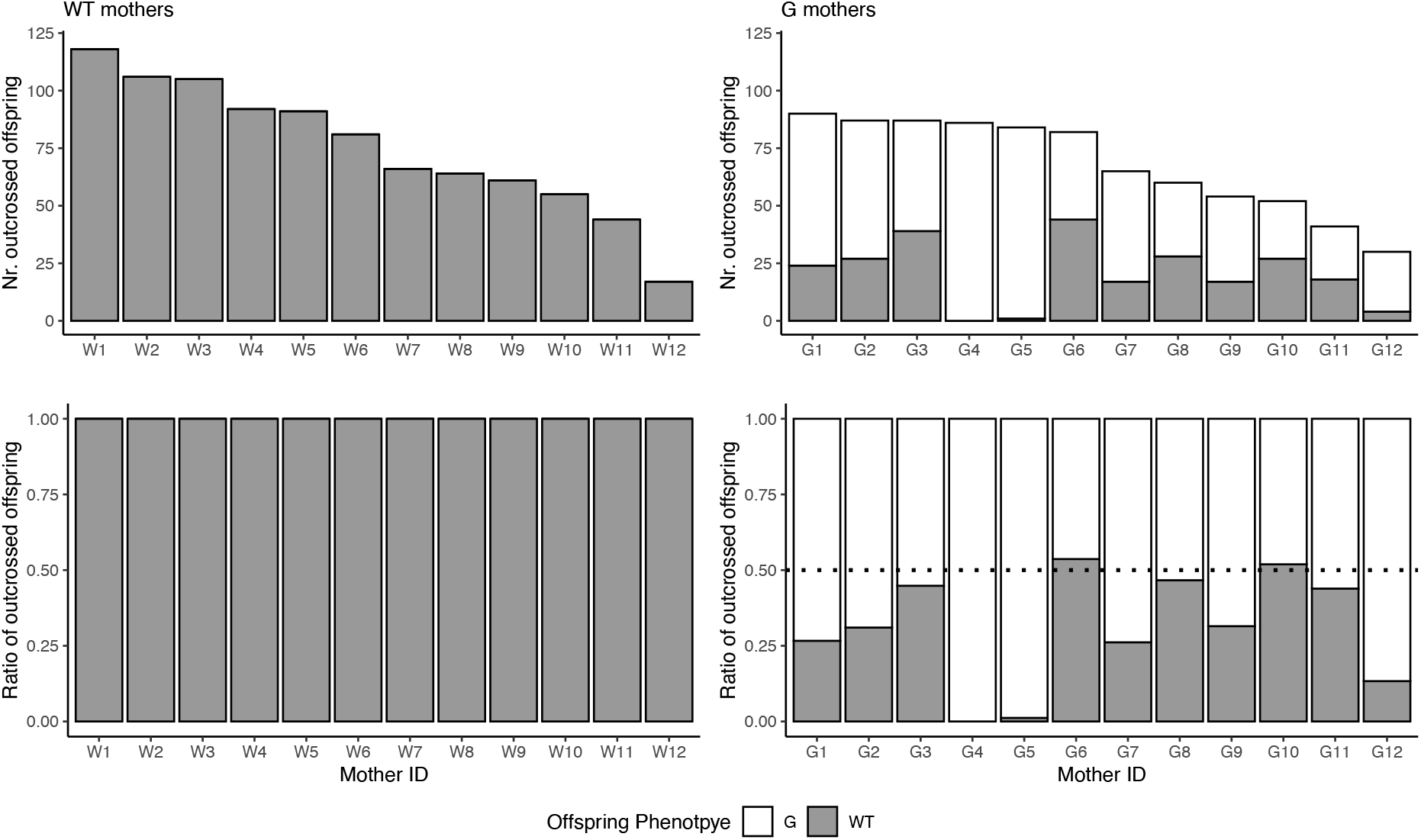
The phenotype of outcrossed offspring from WT and G mothers. The top panels show the number offspring, and the bottom ones show their ratios. The mothers are ordered by the total number of offspring, hence Mother ID does not correspond to Fig. 2. The dotted line in the right bottom panel indicates the expected ratio of segregation or single-locus Mendelian inheritance from heterozygous mothers.

## Discussion

We found that the inheritance mode of Ginger phenotype fits single locus Mendelian inheritance with the dominance of Ginger phenotype over WT. In addition, we did not find any life history difference between G and WT individuals from the same population. Below, we discuss the inheritance of the Ginger phenotype, the potential adaptive significance and future research directions that this finding stimulates.

The phenotypic ratios of selfed and outcrossed offspring suggest that the Ginger shell trait is determined by a single locus, and that the Ginger allele is dominant. We observed that the ratio of selfed G offspring produced by heterozygous G mothers was approximately 0.75 (Fig. 2), and that the ratio of outcrossed G offspring by G mothers became significantly lower than 0.75, but still higher than 0.5 (Fig. 3). The former segregation rate and the change of segregation rate supports single-locus inheritance, while the latter does not perfectly fit to the prediction (see also Fig. S2). This discrepancy could be due to a smaller sample size of outcrossed offspring, as well as to biological reasons. For example, even though we observed that the Greek snails copulated with the lab snails, it is likely that the Greek snails did not use the received sperm for outcrossing but continue (partially) selfing. Previous studies support that *L. stagnalis* prefers outcrossing (Cain 1965; Nakadera et al. 2014), but this is only confirmed in the case when mating partners are from the same population. In addition, we cannot exclude the possibility that G and WT offspring have different hatching success, as we could only score their phenotypes when they are two-month-old. Nonetheless, at this moment, the most likely mode of inheritance of the Ginger phenotype is that the trait is dominant with single-locus inheritance, as found in other model species (e.g., van’t Hof et al. 2016; Feiner et al. 2022; Luttikhuizen and Drent 2008; Kellenberger et al. 2019).

The fascinating question is whether Ginger snails have any ecological advantages in the field. Snails do not see color, so it is unlikely that the trait is used in mate choice as it is in other animal groups (e.g., Cardoso and Mota 2022). Although it is possible that they do not have any advantages (Williams, 2017), two hypotheses come to mind. As mentioned above, their shell color is hardly visible in the adult snails of natural populations due to e.g., algae growing on the shells (Fig. 1A). Thus, if Ginger snails have any advantages directly from their shell color, it would likely be in their juvenile stage when they are building their shells relatively quickly (thus limiting algal growth). Also, we like to point out that their habitat is a relatively shallow lake. So, it is possible that their red color is adaptive against UV radiation or elevated temperature (Han et al. 2022). Moreover, if our speculations are correct and given the difficulties of detecting the phenotype directly in the field, it is plausible that Ginger snails occur in other locations without having been noticed. It would be an interesting follow-up project to make an inventory of field-collected shells as well as museum collection material.

We consider that this newly discovered shell color polymorphism in *L. stagnalis* can provide the opportunities to expand the understanding of the genomic and physiological mechanism to determine shell color as well as the evolution of this trait. This species is fecund, promiscuous, and very easy to maintain in the lab. We also observed that these Greek snails do mate with our lab strain (even though they are same species, it cannot be taken for granted in field collected individuals), thus allowing us to backcross the trait. Finally, compared to *Cepaea*, the shell color polymorphism of *L. stagnalis* is much more straightforward to score. Given the above, we are convinced that the ginger snails provide a great new model system for investigating shell color polymorphism.

In sum, we found a new shell color polymorphism in *L. stagnalis*, and revealed that the Ginger allele is dominant and inherits in a single locus manner. We do not yet know if and why this polymorphism is maintained in the field, since we did not detect any advantage or disadvantage of being Ginger. Lastly, we consider these Ginger snails to be a useful model system to expand the understanding of shell color polymorphism in gastropods.

## Acknowledgement

We appreciate Dr. Marianthi Hatziioannou and Dr. Theodoros Naziridis for helping with collection of snails, Omar Bellaoui and Xaver Bartels for taking care of the snails, and Cathy Levesque for the picture in Figure 1B. This research was supported by Netherlands Organization for Scientific research (NWO) Open Competition Klein to JMK and YN.

