## Supplemental Figures for "Dominant gingers – discovery and inheritance of a new shell polymorphism in the great pond snail *Lymnaea stagnalis*"

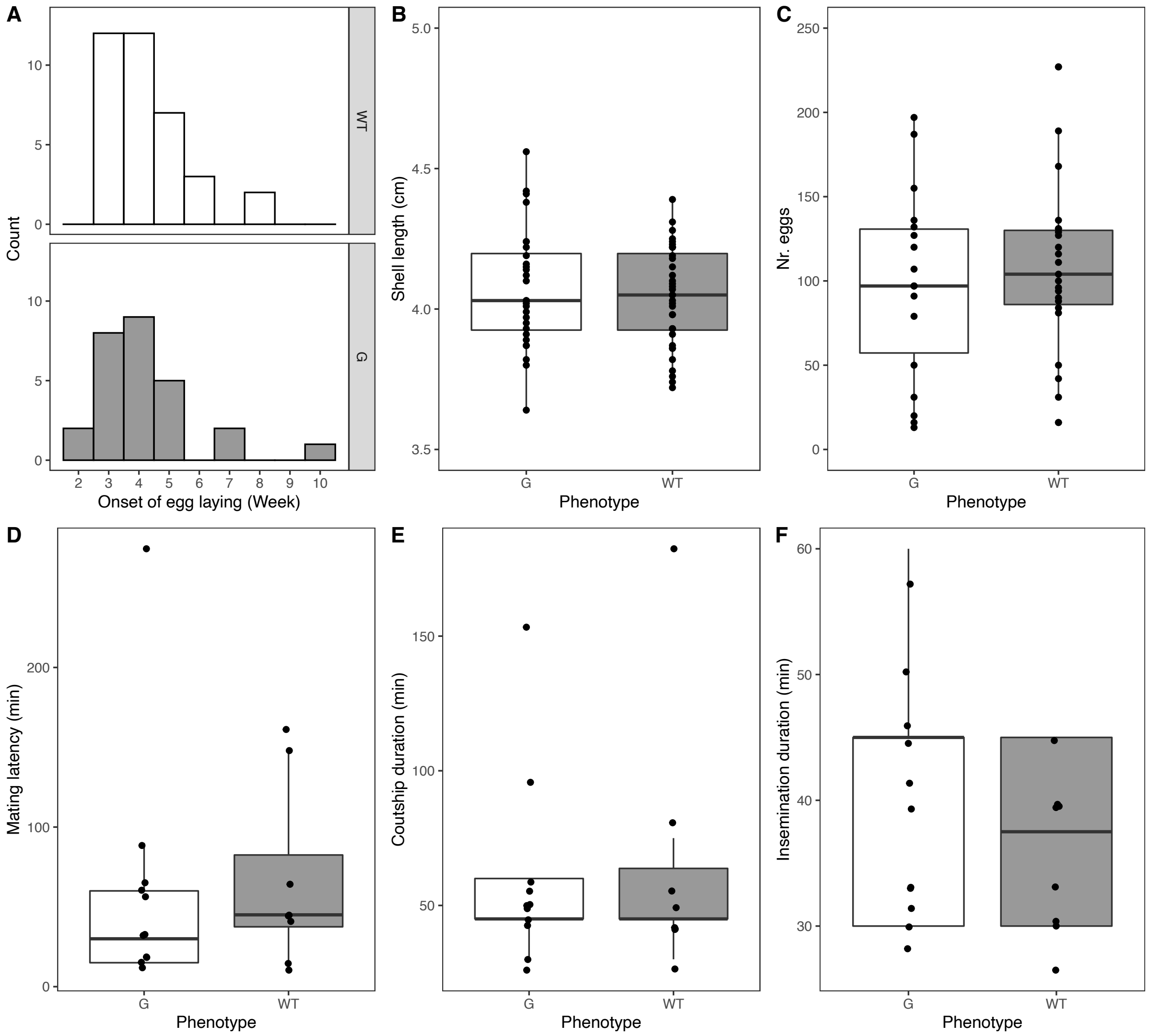


Fig. S1. The comparison of life history traits between Ginger and WT from the Greek population. We measured the onset of egg laying (A), body size (B), the number of eggs produced (C) and mating behaviors (D-F). For the onset of egg laying, one G did not start laying eggs within our 10-week monitoring (N: G = 27, WT = 36), and there was no difference in the onset of egg laying between G and WT ($t_{47.78}$ = -0.16, $P$ = 0.869). Also, at Week 7, we did not detect any difference in body size ($t_{52.16}$ = 0.60, $P$ = 0.548). Moreover, in two days of week 9, we collected 41 egg masses (G = 18, WT = 23) and the number of eggs laid did not differ between G and WT ($t_{34.03}$ = -0.58, $P$ = 0.562). About mating behavior, when the Greek snails mated with the lab snails, 11 of out 12 G and 8 out of 12 WT copulated under our observation, but the mating rates were not different between morphs ($\chi^2_1$ = 2.27, $P$ =

0.132). When they mate, all of the Greek snails acted as female first. In mating pairs, we did not see any difference between morphs in the duration of mating latency ($W$ = 38.5, $P$ = 0.673, removing the outliner did not change the outcome [$W$ = 30.5, $P$ = 0.416]), courtship ($W$ = 41, $P$ = 0.827) or insemination ($W$ = 48, $P$ = 0.745).

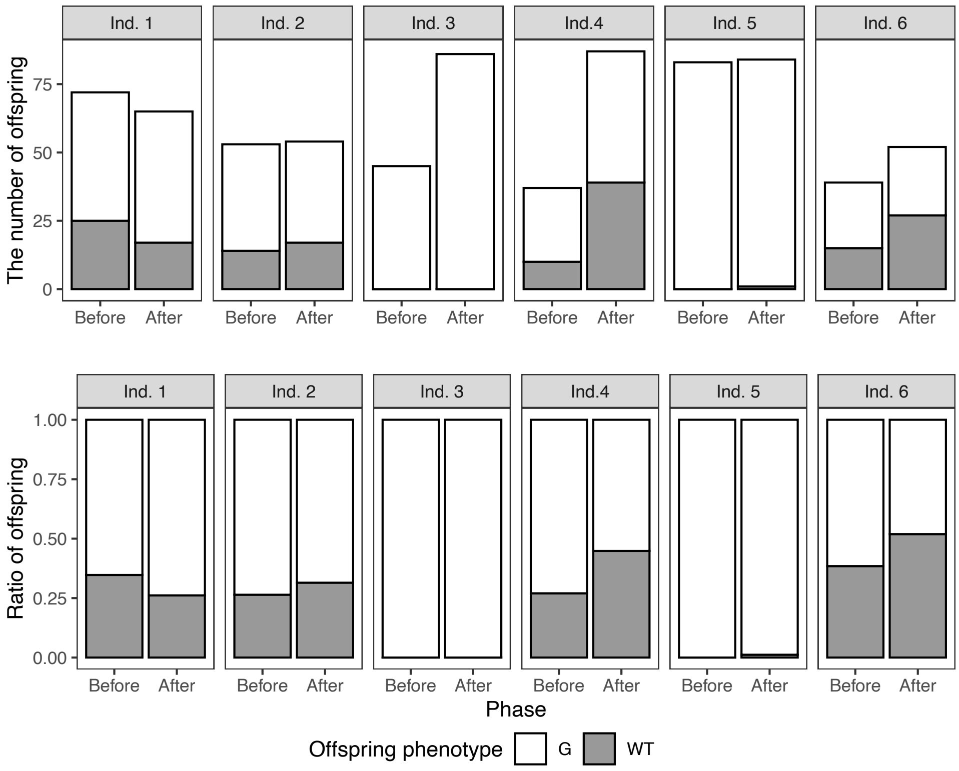


Fig. S2. The phenotype of offspring from G mothers before and after mating. The graphs on the top row show the number of offspring, and those on the bottom row show their ratios. Mother ID corresponds as followed: Ind. 1 = g4, G7; Ind. 2 = g5, G9; Ind. 3 = g8, G4; Ind. 4 = g12, G3; Ind. 5 = g1, G5; Ind. 6 = g10, G10.
